# Transcription terminators with context-dependent promoter and terminator activities

**DOI:** 10.64898/2026.02.12.705533

**Authors:** Letian Bao, Anthony C. Forster

**Affiliations:** Department of Cell and Molecular Biology, Uppsala University, Husargatan 3, Box 596, Uppsala 75124, Sweden

## Abstract

A dogma of RNA synthesis is that promoter activity and termination are unrelated processes governed by completely different DNA sequences. Serendipitously, we find that two class II terminators for bacteriophage T7 RNA polymerase (RNAP), the natural T7 concatemer junction (CJ) and the artificial vesicular stomatitis virus (VSV) terminator, are context-dependent strong *promoters* for *E. coli* RNAP. However, the third class II member, the artificial human preproparathyroid hormone (PTH) terminator, is not a promoter. Transcription start sites and mutagenesis for CJ and VSV reveal a σ^70^ extended TGn promoter motif that is lacking in PTH. Furthermore, results resolve the prior paradox of class II termination apparently occurring only *in vitro*, not *in vivo*, enabling demonstration of class II termination *in vivo* by both T7 and *E. coli* RNAPs. In addition, engineering of T7 RNAP increases its efficiency of class II termination *in vivo*. Given that prokaryotic synthetic biology is reliant on class I (hairpin) terminators, with T_Φ_ used almost exclusively in constructs for T7 RNAP, recombination is an issue in large constructs. Thus, small class II terminators, now better understood with respect to terminator and promoter activities in the context of various sequences and RNAPs, extends the synthetic biology toolkit.

## INTRODUCTION

Bacteriophage T7 RNA polymerase (T7 RNAP) is widely used in molecular biology but there are only two natural terminators of its RNA synthesis (Studier *et al*., 1990; Borkotoky and Murali, 2018). T_Φ_ is a class I, intrinsic terminator which encodes a hairpin followed by a short stretch of uridines (Dunn and Studier, 1983). Transcription termination is only 80% efficient as leakage is needed for low-level expression of downstream genes lacking their own promoters (Macdonald *et al*., 1994). The concatemer junction (CJ) pauser/terminator is a class II, shorter sequence (Lyakhov *et al*., 1997). Homologous unnatural class II termination sequences were discovered by accident within the human preproparathyroid hormone gene (PTH, Mead *et al*., 1986) and vesicular stomatitis virus cDNA (VSV, Whelan *et al*., 1995). These three sequences are 15-18 base pairs and share a seven-base strictly conserved motif that must be preceded immediately by A, C or T for pausing/termination (He *et al*., 1998; Fig. 1A). Unlike class I, class II are unlikely to terminate via secondary structure formation. Instead, the mechanism is apparently via destabilization of the DNA:RNA hybrid compared to the competing DNA double helix at the seven-base conserved site (Zhang and Russu, 2014; Fig. S1). Because of their small size, we proposed that class II terminators might be ideal for multigene constructions in synthetic biology by minimizing potential recombination between repeats of the sequences (Bzymek and Lovett, 2001). However, in contrast to efficient termination *in vitro* (∼70% for VSV, Lyakhov *et al*., 1998), the VSV and PTH terminations appeared to be severely inhibited in *E. coli* cells for an unknown reason (Du *et al*., 2009, 2012). This intriguing paradox motivated us to test ever more constructs for T7 RNAP, which led us to study promoters.

**Figure 1.**
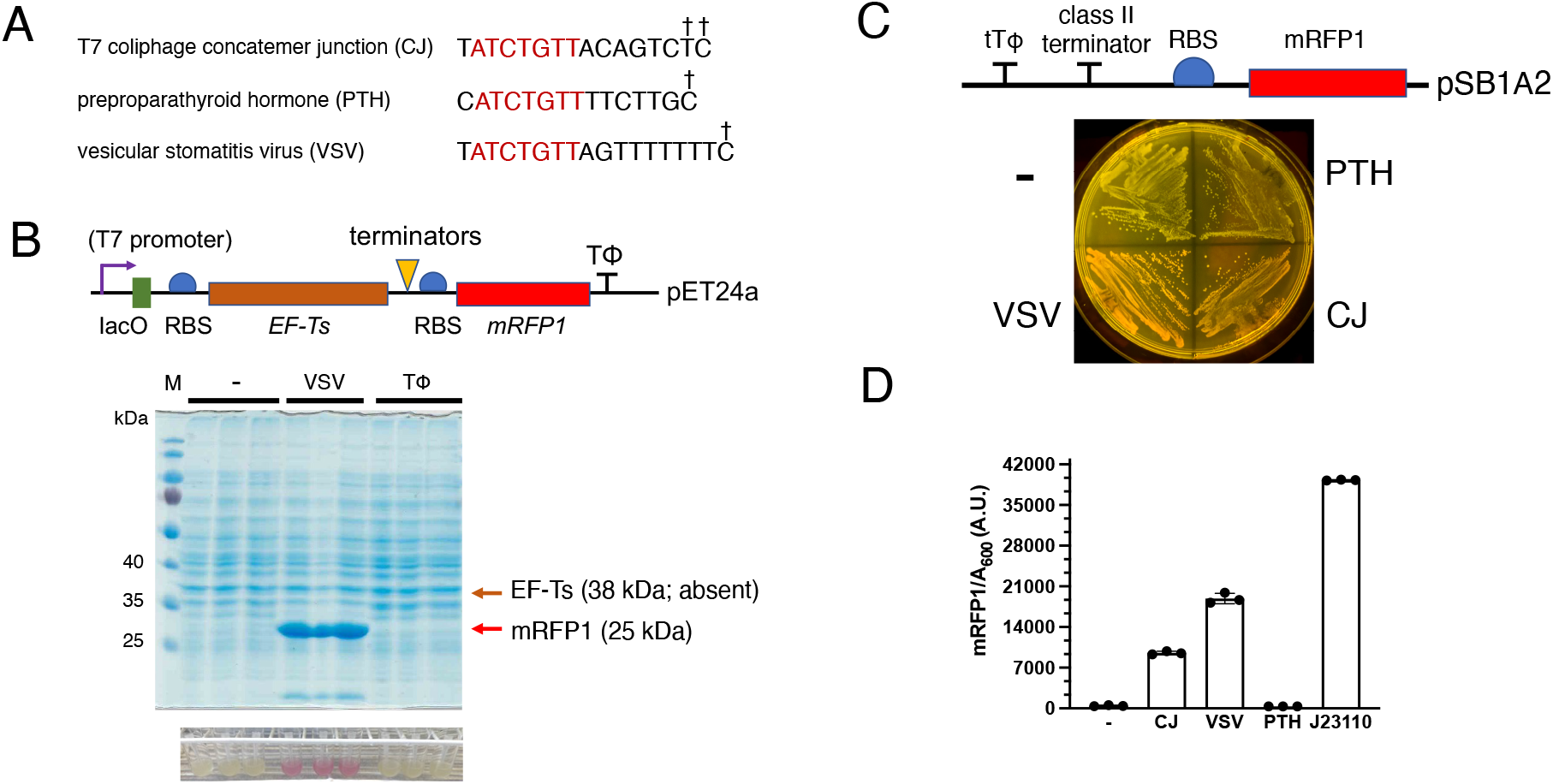
Discovery of promoter activities for *E. coli* RNAP associated with two terminators for T7 RNAP. (A) Sequence alignment of the three class II terminators for T7 RNAP. Red, seven-base strictly conserved sequence; †, termination site (Lyakhov *et al*., 1998). (B) Constitutive expression of mRFP1 from bicistronic constructs in *uninduced* overnight liquid cultures of *E. coli* DH5α visualized in ambient light and by Coomassie Blue staining of SDS-PAGE (triplicates). Note that IPTG induction controls produced additional strong bands corresponding to EF-Ts (not shown). (C) Fluorescence of re-streaked *E. coli* colonies of monomeric constructs visualized through an orange screen upon irradiation with blue light. (D) Quantitation of mRFP1 expression in overnight liquid cultures of *E. coli* transformed with the monomeric constructs of (C). Error bars are standard deviations, n=3.

T7 RNAP initiates transcription from a highly-conserved T7 promoter, which is orthogonal to *E. coli* promoters (Basu and Maitra, 1986). In contrast, a typical *E. coli* promoter consists of two consensus hexamer sequences: the -10 element (or Pribnow box) and -35 element, named according to their location upstream of the transcription start site (Harley and Reynolds, 1987). These motifs serve as binding sites for RNA polymerase holoenzyme, which consists of the core enzyme and a σ factor. *E. coli* encodes seven σ factors and each directs RNA polymerase core enzyme to specific promoters under different conditions (Tab. S1; Ishihama, 2000; Gruber and Gross, 2003; Paget, 2015).

Structural studies have revealed the molecular basis of promoter recognition by σ factors. At the -10 element, the region 2 of the σ factors interact with highly conserved nucleotides, -11A and -7T, promoting base flipping and stabilization within hydrophobic and hydrophilic pockets of the protein, respectively (Feklístov and Darst, 2011). These interactions facilitate promoter melting and stabilize the open complex, supporting a model in which recognition of the -10 element actively drives promoter opening (Feklistov and Darst, 2011). Recognition of the -35 element is mediated by a helix-turn-helix motif within region 4 of the σ factors. Structural analysis revealed specific interactions, including the -35T at the non-template strand, and the - 34A, -33C, -31G at the template strand (Campbell *et al*., 2002), which correlates well with earlier mutational studies demonstrating the importance of region 4 for promoter recognition (Gardella *et al*.,1989; Sharp *et al*., 1999).

Promoter activity is also influenced by sequences located between the −10 and −35 elements. An AT-rich spacer can enhance transcription by increasing RNA polymerase binding and facilitating open-complex formation (Liu *et al*., 2004; Hook-Barnard and Hinton, 2009; Klein *et al*., 2021). Genome-wide alignment of *E. coli* promoters also revealed a preference for Ts at positions -17 and -18 (Mitchell *et al*., 2003), which are recognized by the R451 of the σ^70^ to increase promoter strength (Singh *et al*., 2011). Indeed, many intragenic promoters depend on an AT-tract located between positions -17 and -23, enabling DNA bending and interaction of the backbone with R451 (Warman *et al*., 2020).

In addition to many promoters containing both -35 and -10 motifs, some promoters work without a -35 element, relying instead on an extended -10 motif, 5′-TGn-3′, which is recognized by the region 2.5 of the σ^70^ factor to compensate (Gralla, 1996, Barne *et al*., 1997). Conversely, others lack a good -10 element but remain active when supported by an extended -10 motif and a -35 element (e.g. bacteriophage T4 P_minor_ promoter; Hook-Barnard *et al*., 2006). Such variability complicates promoter predictions. In *E. coli*, in addition to genome-wide promoter mapping (Urtecho *et al*., 2023), recent studies combining systematic functional assays and computational analysis provide mechanistic insights into promoter recognition (John *et al*., 2022). Furthermore, developments in promoter prediction tools based on machine and deep learning have improved discrimination between promoter and non-promoter sequences (Paul *et al*., 2024). A few reports experimentally identified promoter elements within random sequences (Wolf *et al*., 2015; Yona *et al*., 2018; Lagator *et al*., 2022; Fuqua and Wagner, 2026). However, these reports focused on evolutionary questions and homologies with typical σ^70^ promoters rather identifying other promoter elements (see below).

## RESULTS

### Insertions of two of three class II terminators for T7 RNAP promote efficient *transcription* downstream by *E. coli* RNA polymerase

Bicistronic reporter plasmids were constructed (Fig. 1B top) to assay termination *in E. coli* by class II terminators inserted between two protein-coding regions. For the *uninduced* negative controls, the expectation was that neither of the two proteins would be produced. Yet, surprisingly, strong red coloration of the overnight cultures was seen (Fig. 1B bottom) from the mRFP1 downstream gene when the VSV terminator was inserted directly upstream. Very strong expression of mRFP1 in the absence of expression of the upstream gene (EF-Ts) was confirmed on a gel (Fig. 1B). However, expression was absent when the VSV sequence was absent (Fig. 1B). This suggested intrinsic promoter activity due to the VSV terminator rather than a cryptic promoter within the EF-Ts gene. To minimize potential background expression from transcripts originating elsewhere on the plasmids (Neff and Chamberlin, 1980), we next substituted the T7 promoter and EF-Ts gene with a truncated native T_Φ_ terminator (tT_Φ_, Calvopina-Chavez *et al*., 2022) and also tested all three class II terminators (Fig. 1C top). Fluorescence in agar (Fig. 1C bottom) and liquid cultures showed promoter activity due to the VSV terminator at 48% of the standard strong BioBrick promoter J23110, while the CJ terminator gave 25% (Fig. 1D). In contrast, the PTH terminator lacked promoter activity relative to promoter-less controls. This implied that the conserved seven-base termination sequence alone was insufficient for promoter activity.

The *in vivo* transcription start sites (TSSs) due to CJ and VSV were mapped by primer extension (Fig. 2) to just a few bases downstream of the termination bases (Fig. 1A), further supporting importance of the terminator sequences in promoter activity. As a positive control, the TSS of the J23110 promoter was located at +2A (Fig. S2 left), presumably reflecting the purine initiation preference of *E. coli* RNAP (Walker and Osuna, 2002; Vvedenskaya *et al*., 2016).

**Figure 2.**
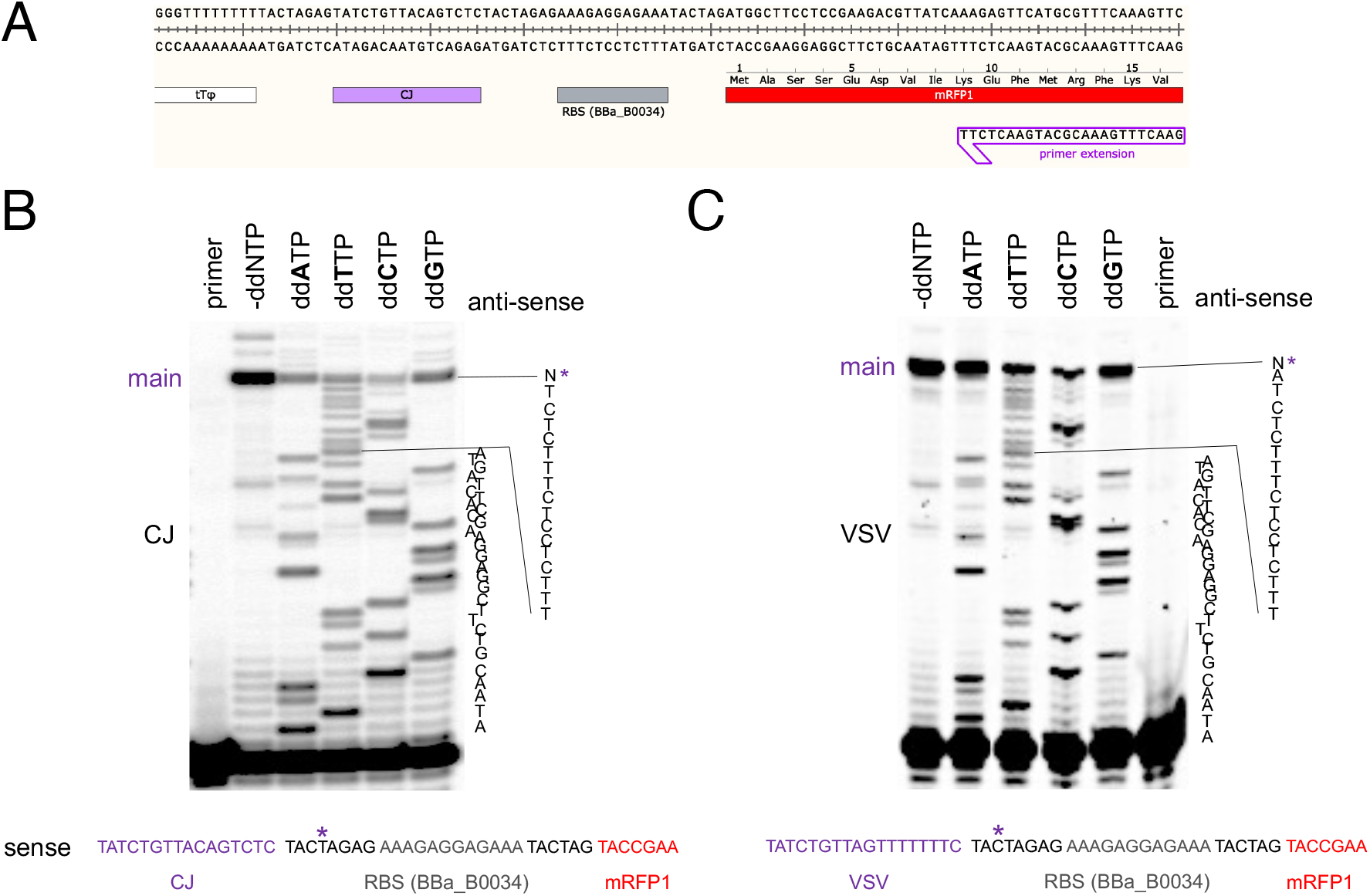
Mapping of *in vivo* transcription start sites of CJ and VSV promoter constructs. (A) Sequence of CJ monomeric plasmid and bound primer. Representative gels of CJ (B) and VSV (C) visualized by phosphorimaging. Main stop sites for reverse transcriptase are marked with asterisks. The whole gels are shown in Fig. S3.

### Identifying our promoter motif

Three truncation mutants were constructed within the VSV terminator sequence of the monomeric construct (Fig. 3A) and their promoter activities were compared at three different copy numbers (Fig. 3B-D). Compared with controls, VSV middle and polyTs alone had little activity. Rather, major promoter activity correlated with the segment containing the entire conserved termination sequence. Copy number affected relative activities quantitatively in a couple of cases, but it did not affect the overall trends greatly. As the high-copy assays were most sensitive and straightforward, we continued with those to further define the promoter motif.

**Figure 3.**
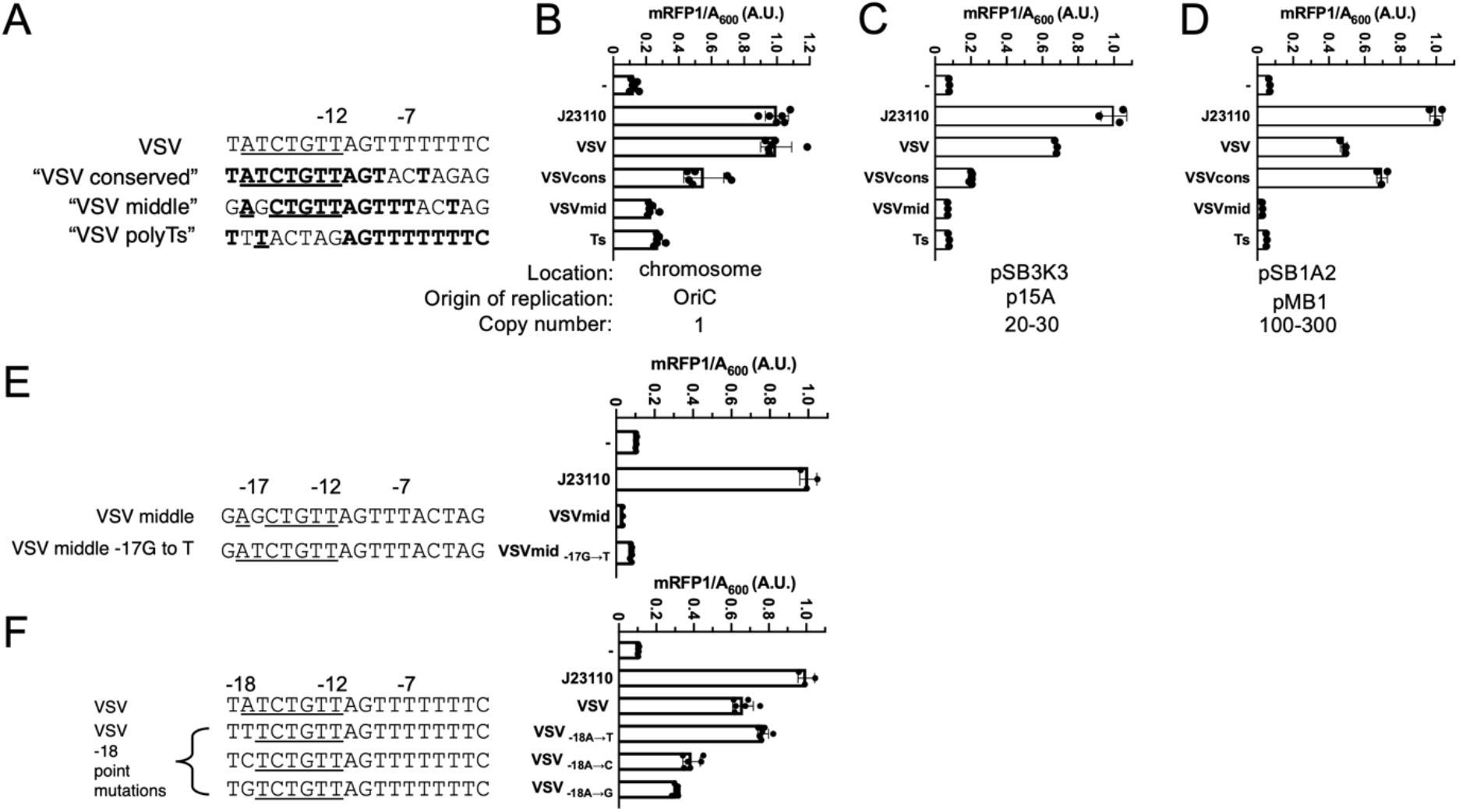
Identifying the role of the VSV terminator sequence in promoter activity. (A) Sequence alignment of truncated VSV terminator sequences in the monomeric construct. mRFP1 fluorescence was measured of constructs on the *E. coli* chromosome (B), a medium-copy plasmid (C) or high-copy plasmid (D). (E) -17G to T point mutation on VSVmiddle to restore the conserved sequence. (F) -18A point mutations to disrupt the conserved sequence. Conserved sequences in class II terminators are underlined, and homologous sequences with VSV in (A) are in bold. All the values are normalized to J23110. Error bars are standard deviation, n≥3.

Next, the -17G of VSVmiddle was mutated to T to restore the entire conserved termination sequence (Fig. 3E). Though an increase in relative activity was seen, the total activity was similar to the negative control (minus class II terminator). Also, all three mutations of the -18A of VSV that altered the conserved sequence had effects but still gave promoter activities between those of the positive (J23110) and negative controls (Fig. 3F). Together, these results indicate that the conserved termination sequence is important, but neither sufficient (see also Fig. 1C, D) nor completely required for promoter activity. In other words, outside sequences were also implicated.

Knowledge of the TSSs of the class II terminator promoters aided alignment with the J23110 promoter and the two σ^70^ consensus promoters to search for homologies outside the conserved termination sequence (Fig. 4A). Alignments slightly shifted from the one shown are possible (Mazumder and Kapanidis, 2019) but they decrease homologies. The VSV bicistronic construct (Fig. 1B) is also aligned; it exhibited high promoter activity despite sequence divergence nine bases upstream of the conserved seven-base termination sequence. Another sequence aligned is the substitution mutant of tT_Φ_ with T_Φ_ in the monocistronic VSV construct, which retained the same strong promoter activity (Fig. S4). Thus, although the important first dinucleotide - 35TT (Campbell *et al*., 2002) of the typical σ^70^ consensus is shared by some of our strong promoters and J23110, considerable variation in the -35 box region is tolerated. The -10 box region also varies, though all of our strong promoters share the -12TA dinucleotide subset.

**Figure 4.**
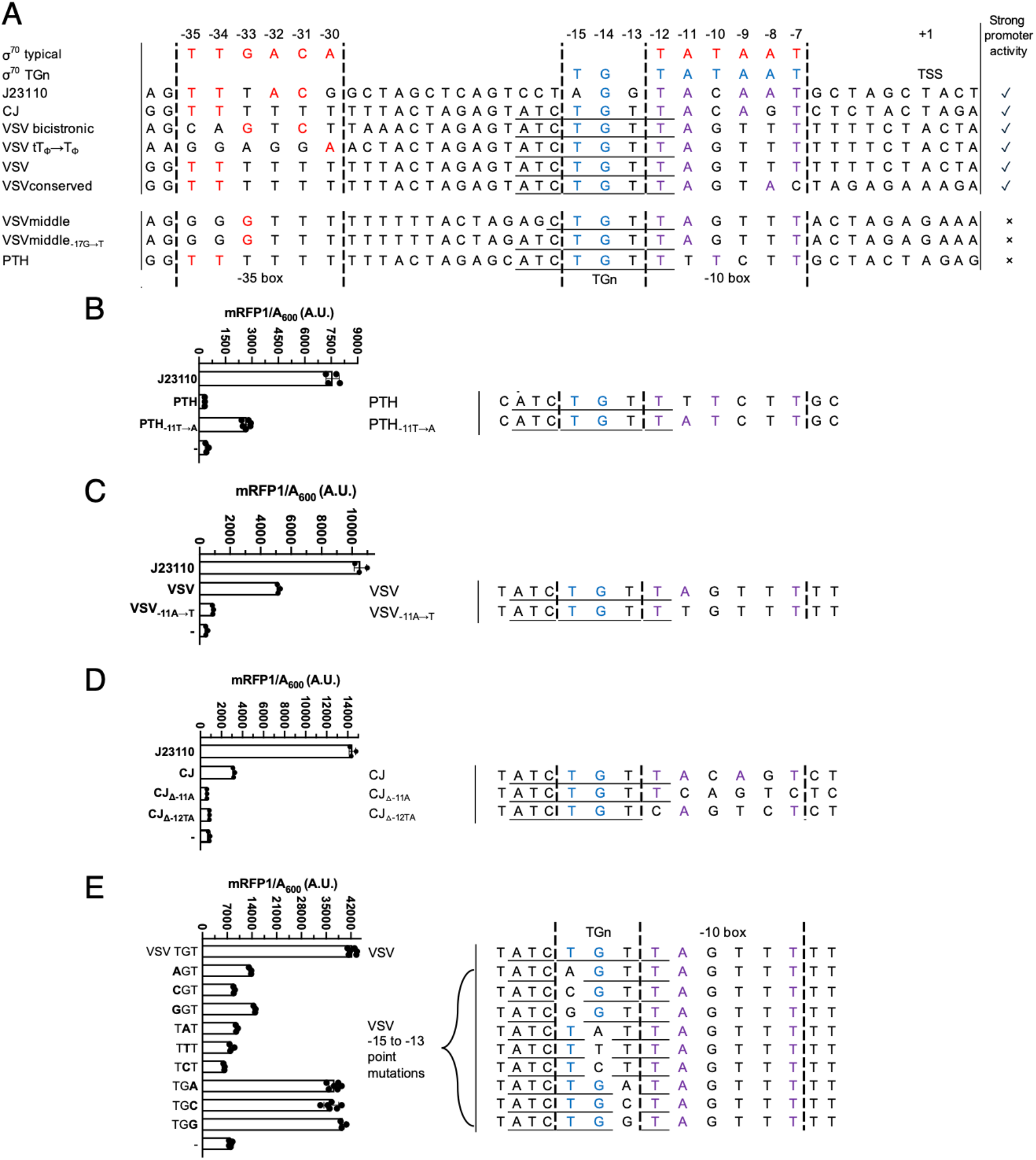
Identifying the role of the larger terminator region in promoter activity. (A) Alignment of terminator constructs with sequences recognized by the σ^70^ transcription factor. Conserved bases of class II terminators are underlined, matches to the σ70 typical consensus are red, to σ70 TGn are blue, and to both are purple. Mutational analysis of PTH (B), VSV (C) and CJ (D) promoters measured by mRFP1 fluorescence. (E) Effects of point mutations in the -15 TGn region on VSV promoter activity. Error bars are standard deviations, n≥3.

As ∼20% of σ^70^ promoters that lack strong -35 box homology are compensated by what is known as the extended -15TGn motif (Fig. 4A) which is also associated with -12T, -11A and -7T (Mitchell *et al*., 2003), we checked for this possibility. Notably, -15TG is shared by all our strong promoters, but also by the poor VSVmiddle and PTH. Having the weakest -35 box in Fig. 4A may explain the poor activity of VSVmiddle and VSVmiddle -17G to T, but not of PTH, which has the same -35 box as VSV in the alignment. This focused our attention back on the -10 box, where PTH stands out as the only sequence lacking -12TA. To test the significance of this correlation, we substituted -11T by A in PTH and -11A by T in VSV (Figs. 4B, C). Indeed, these mutations dramatically increased and decreased the activities, respectively, strongly supporting the hypothesis. Deletion mutagenesis further supported the significance of the -10 box region (Fig. S5), with the importance of -12T revealed in the CJ promoters (Fig. 4D). Furthermore, saturation point mutagenesis of the TGn motif in VSV confirmed the importance of this exact TGn sequence for promoter activity, in good agreement with Mitchell *et al*. (2003) (Fig. 4E).

Returning to the variability of the -35 box, 11 addition mutants were constructed in this region, ranked by activity and examined for homology with the typical σ^70^ -35 box consensus (Fig. S6). Some correlation is seen with the overall number of Ts in the -35 box, although not absolute. Also, we measured promoter activities of 13 randomized spacer sequences between the -35 and -10 boxes (Fig. S7A). This suggested that the AT richness of the sequence between the boxes stimulated promoter activity, consistent with previous observations (Klein *et al*., 2021). Commonalities between the heat map and sequence logo analyses showed that A and T are more preferred than G and C in the spacer (Fig. S7B, C). Next, to gain further insight into the VSV promoter motif, we analyzed promoter activities from large random DNA libraries.

### Analysis of promoter databases

We found three reports of prokaryotic promoter elements identified from random sequence libraries. Yona *et al*. (2018) replaced the native lac promoter with various 103-base randomized sequences (each is equivalent to 73 overlapping randomized 30-mers) upstream of a modified *lac* operon in the *E. coli* genome. Only cells carrying functional promoter sequences could utilize lactose and thus grow on M9–lactose plates. This assay, although not a quantitative readout of promoter strength, enabled distinguishing active from essentially inactive promoters. Out of 40 randomized sequences, only four supported growth, all of which were attributed to typical σ^70^ binding sites. In subsequent evolution experiments, single point mutations were sufficient to activate every randomized sequence, also attributed to creating typical σ^70^ binding sites. We searched their 36 essentially inactive promoters (free of a typical σ^70^ binding site) for the major contiguous segment of our VSV promoter, -15 TGTTA (Fig. 4A). We found it in seven (19%), confirming that sequences outside the TGTTA are needed for promoter activity (as seen above with VSVmiddle).

Lagator *et al*. (2022) constructed a diverse library of 13,203 sequences by fully randomizing a 36-nucleotide region (*36N* library; 25% mutation rate per position) cloned upstream of a yellow fluorescent protein (YFP) reporter on plasmids, transformed it into *E. coli*, and assayed it using fluorescence-activated cell sorting (FACS). The library exhibited a 26.7% probability of members having fluorescence and contained a significant number of active promoters lacking the typical σ^70^ binding site but containing TGTTA (116, Fig. 5A, left). Converting the data into a heatmap revealed a putative −35 element TTAGGG (Fig. 5A, right). Three additional regions of homology were detected between the VSV promoter and the *36N* sequences: - 40 AGG, −35 TT and -19 TAT (Fig. 5A, right). This supports potential roles of specific sequences upstream of the TGTTA in promoter activity.

**Figure 5.**
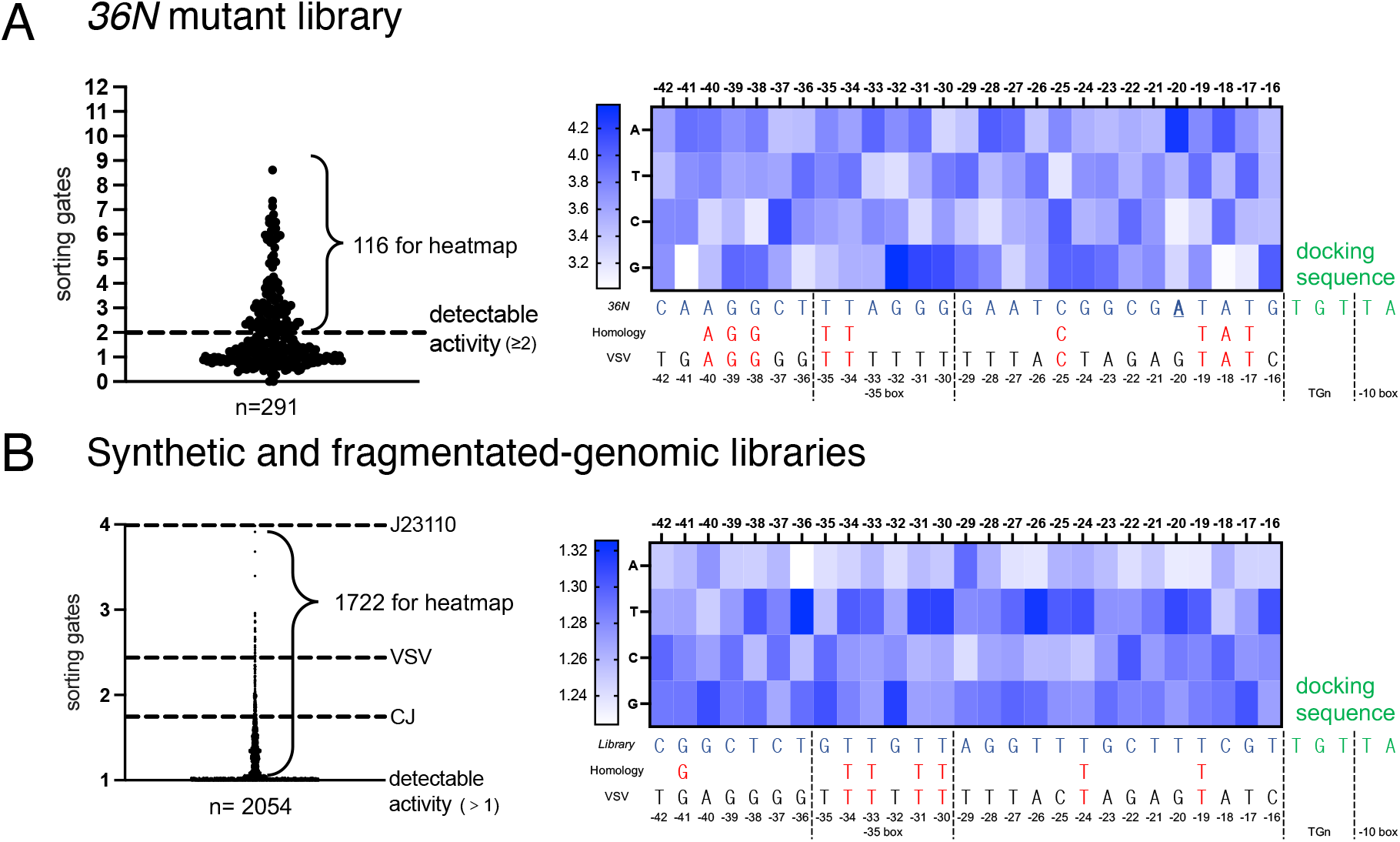
Analysis of Lagator *et al*. (2022) and Fuqua and Wagner (2026) datasets for promoters lacking the typical σ^70^ binding site but containing TGTTA. (A) Analysis of the fully-randomized *36N* promoter library in Lagator *et al*. (2022). Of 291 sequences lacking the typical σ^70^ binding site but containing TGTTA, 116 had detectable activity. The heatmap summarizes these 116 sequences, with rows representing individual bases and columns representing base positions upstream of the motif. Heatmap color intensity reflects the correlation between base at each position and relative promoter activity, with darker blue indicating bases preferentially enriched in higher-activity promoters and lighter shades indicating enrichment in lower-activity promoters. (B) Analysis of Fuqua and Wagner (2026) datasets containing 2,054 sequences lacking the typical σ^70^ binding site but including the TGTTA motif, of which 1,722 exhibited detectable promoter activity (left). Our VSV and CJ activities are compared with the data by using the common J23110 standard. The heatmap of these 1,722 (right) was plotted as in (A). Red bases indicate homology between the bases from the library correlating with high activity and the VSV sequence. Sequences were classified as active promoters when their sorting gate was equal or greater than 2 in (A), and greater than 1 in (B), and sequences with a sorting gate equal to 1 were classified as undetectable activity. For the library in (B), an alternative match to the TTGACA -35 box is achieved by reducing the spacer by 1 base.

Our most meaningful analysis, due to the library size and internal control, was performed using the dataset of Fuqua and Wagner (2026). This comprised 17,129 synthesized random 150-bp sequences and 91,866 fragmented *E. coli* genomic sequences averaging 130 bp. These libraries were cloned into a bidirectional fluorescence reporter plasmid such that promoters on the top and bottom strands drove GFP and RFP expression, respectively. BBa_J23110 was their positive control and standard for calculating relative promoter activities by FACS. We identified a substantially larger number of promoters lacking the typical σ^70^ binding site but containing the TGTTA, including 959 top-strand promoters (574 synthetic + 385 genomic) and 763 bottom-strand promoters (532 synthetic + 231 genomic) = 1722 of a total of 2054 sequences (Fig. 5B, left). Because BBa_J23110 was used as the common reference in both our experiments and those of Fuqua and Wagner (2026), we could directly calibrate our promoter activities of the CJ and VSV promoters with activities of their library members. Among the 1,722 promoters lacking the typical σ^70^ binding site but containing the TGTTA, only 176 exhibited activities ≥ CJ and 34 > VSV, indicating that both CJ and VSV are among the strongest. Heatmap analysis identified a promoter consensus where the four Ts in the -35 GTTGTT accounted for the contiguous homology with the VSV promoter (Fig. 5B, right). The lack of consensus in the -16 to -18 region supports the results of Mitchell *et al*. (2003) and our conclusion above that the poor activity of VSVmiddle was due to a weak -35 box rather than an altered -17 base.

A genome-wide search of *E. coli* MG1655 (GenBank: U00096.3) identified 30,944 instances of the TGnTA motif. Combining with the global TSS mapping dataset from Thomason *et al*. (2015), we focused on the 13,247 TSSs not overlapping with annotated RegulonDB sites (*i*.*e*. essentially not overlapping with σ-binding promoters upstream of genes; Saldago *et al*., 2013). Interestingly, 833 (6.3%) harbored a TGnTA motif precisely at −15 positions and 2,043 (15.4%) contained the motif within a ±5 base shift. Among their 492 “orphan” subset, 30 (6.1%) contained TGnTA at −15 position and 67 (13.6%) within a ±5 bp shift; sequence logo analysis indicated a slight favoring of Ts across the promoter region (Fig. S8). These data indicate that promoters similar to ours may be important for widespread transcriptional expression and regulation in *E. coli*.

### Termination *in vivo* of T7 and *E. coli* RNAPs by class II terminators

Finally, the discovery of cryptic VSV promoter activity led us to solve a prior paradox in class II T7 termination. Almost all work with these terminators had been done *in vitro*. Yet, when we attempted reproduction *in vivo* using the strongest class II terminator, VSV, the most straightforward interpretation of our puzzling data was that the terminator was very inefficient *in vivo* (Du *et al*., 2009, 2012; Fig. S9, top). Here, we propose that VSV did terminate efficiently, but this was masked by its strong cryptic promoter activity (Fig. S9, bottom). To circumvent such confounding promoter activity, we made new bicistronic constructs where the −35 element of class II terminators was predicted to be poor (CACCGG of Mut 7 in Fig. S6) and the confounding lacO operator upstream of the second gene (Fig. S9) was removed (Fig. 6A). Indeed, fwYellow synthesis was negligible in uninduced controls (Fig. S10), revealing minimal, if any, cryptic promoter activities. Induction of transcription by T7 RNAP now unmasked *in vivo* termination activities of all three class II terminators placed between the two fluorescent genes, though tTφ was the strongest (Fig. S10; Fig. 6B). Having established *in vivo* termination of T7 RNAP by class II terminators, we wondered whether they could also terminate *E. coli* RNAP. Indeed, replacing the T7 promoter with the constitutive *E. coli* promoter BBa_J23110 resulted in similar termination efficiencies (Fig. 6C and D. Thus, in the appropriate sequence context, T7 class II terminators can act as promoters or terminators *in vivo* of the very same enzyme, *E. coli* RNAP.

**Figure 6.**
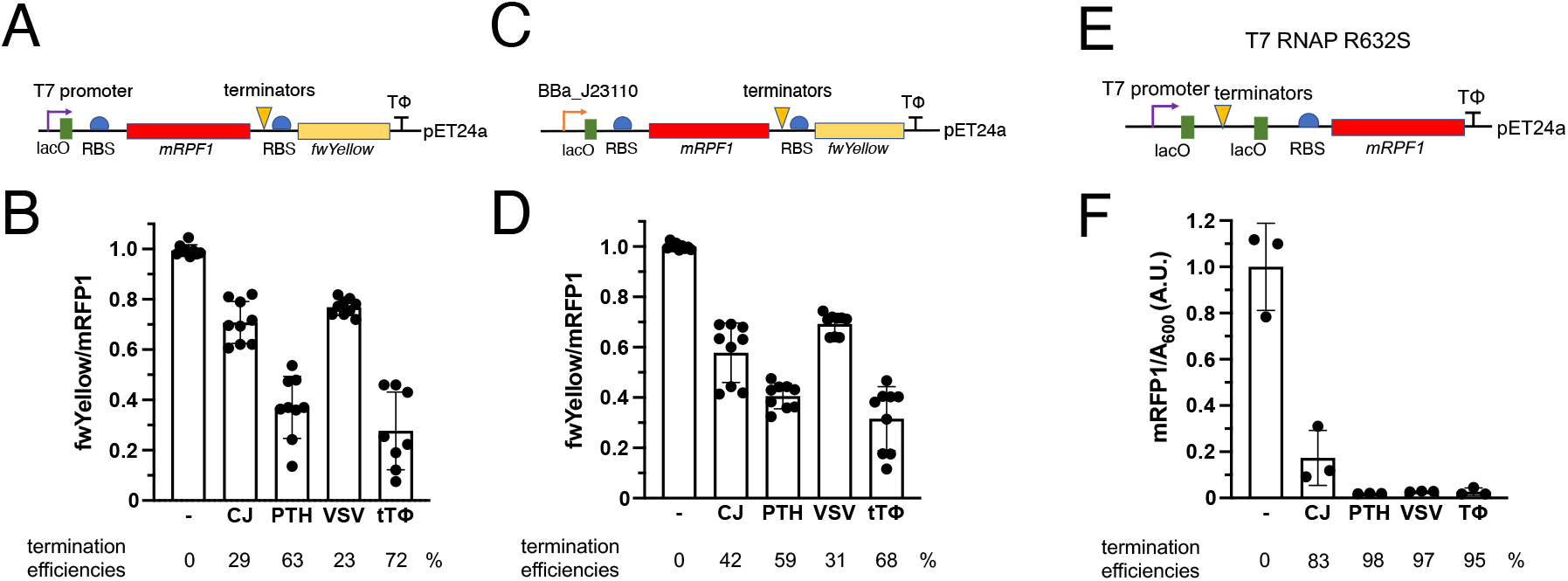
Termination efficiencies *in vivo* of class II T7 RNAP terminators on T7, *E. coli* and mutant T7 RNA polymerases. (A) Schematic of the redesigned bicistronic construct used to assay termination efficiency of T7 RNAP. Terminators were placed between mRFP1 and fwYellow. (B) Quantification of termination efficiencies on T7 RNAP based on fwYellow/mRFP1 ratios measured by fluorescence following 4 h 1 mM IPTG induction. (C) Schematic of the bicistronic construct used to assay termination efficiency on *E. coli* RNAP, in which the T7 promoter was replaced with the constitutive BBa_J23110 promoter. (D) Quantification of termination efficiencies on *E. coli* RNAP based on fwYellow/mRFP1 ratios measured by fluorescence after 24 h 10 mM IPTG induction in *E. coli* DH5α. Ratios were normalized to the construct lacking an intervening terminator. (E) Schematic of the monocistronic reporter and fluorescence quantification of mRFP1 expression downstream of different terminators. (F) Fluorescence-based termination efficiency analysis of class II terminators in monocistronic constructs transcribed by R632S T7 RNAP. For a comparison with wild-type enzyme, see Fig. S10. Expression levels were normalized to the construct lacking a terminator. N≥3, error bars are standard deviations.

The remaining challenge to facilitate applications of class II terminators in synthetic biology was to how to achieve terminations as efficient as achieved by the class I terminator Tφ? One approach is to use these short sequences in tandem (Du *et al*., 2009, 2012). A second potential approach evaluated here is to use a mutant T7 RNAP. As a termination study by Temme *et al*. (2021) used T7 RNAP with a mutation (R632S) in a region previously associated with polymerase processivity, we wondered if it might give different termination efficiencies compared with the wild-type enzyme. Thus, we constructed this mutant gene on the *E. coli* MG1655 chromosome and indeed found that the mutant had dramatically increased termination efficiencies (Fig. 6E and F), presumably due to decreased processivity.

## DISCUSSION

We were surprised to find that suitably placed terminators can be strong promoters after so many decades of molecular biology and initially wondered if overlapping terminator-promoter sequences made sense, mechanistically. Superficially, both promoter activity and class II termination involve melting of short double-stranded nucleic acid helices. The promoter -10 box is AT rich to aid DNA melting by the σ factors and it is positioned at a similar distance from the TSS as the termination sequences (Fig. 4A) which also favor melting, but of DNA-RNA hybrids. However, the conserved termination sequence is apparently not terminating simply by having the lowest possible stability of the DNA-RNA hybrid, but rather by the hybrid having a lower relative stability than the competing DNA double helix (Zhang and Russu, 2014; Fig. S1). Furthermore, the related PTH version was not a promoter (Fig. 1D) and the polyT portion of the VSV terminator was not key for promoter activity (Fig. 3A). Instead, our mutagenesis showed that the promoter activity derives from overlap with the extended -15 TGnTA motif shared by ∼20% of σ^70^ promoters that lack strong -35 box homology (Mitchell *et al*., 2003) and that activities also correlated with Ts in the -35 box and AT richness of the spacer.

The only natural class II terminator, CJ, is located in the early region of the phage T7 genome that is transcribed by *E. coli* RNA polymerase, 750 bases upstream of the next gene (gene 0.3). So, this unassigned long intragenic sequence of T7 may have a function at the RNA level early in infection before expression of the T7 RNAP. However, stable transcripts from this region were not abundant in transcriptomic studies of early infection (Jack *et al*., 2017). In general, genome-wide studies report pervasive transcription initiation from regions lacking annotated promoters (Thomason *et al*., 2015) that may include the extended -15TGn motif and combined terminator-promoters. Furthermore, a combined terminator-promoter enables a form of recycling where RNAP releases the RNA transcript but remains bound to DNA, diffusing one-dimensionally to nearby promoters to facilitate reinitiation. Recycling has been demonstrated by single-molecule studies with class I hairpin terminators (Song *et al*., 2024) and may not only increase the efficiency of transcription, but also streamline genomes (Fletcher and Grainger, 2024), as many viral, bacterial and plastid genomes have minimal intergenic spacers. Indeed, combined terminator-promoter constructs have recently been created artificially for yeast synthetic biology (Ni *et al*., 2023).

Our results also improve the toolkit for synthetic biology, given that essentially all constructs for expression via T7 RNAP rely on one large terminator, T_Φ_, which is prone to recombination if duplicated in multigene constructs (Shepherd *et al*., 2017). Having now demonstrated that small class II terminators function *in vivo* for T7 and *E. coli* RNAPs and are even more efficient with an engineered T7 RNAP (Fig. 6F), they provide recombination-resistant alternatives. For termination applications, PTH should be favored over CJ and VSV because of the absence of confounding cryptic promoter activity and higher termination efficiency *in vivo*. We thus propose that small terminators and extended TGn promoters, overlapping or not, in the context of T7, mutant T7 or *E. coli* RNAPs, should enhance synthetic biology studies.

## MATERIALS AND METHODS

### Constructions and mutagenesis

Constructions of plasmids for termination assays in *E. coli DH5a* were by BioBrick cloning (Knight, 2003). Amplification of the expressing cassettes (Fig. 1C) for integration into the *Acatsac* region of the chromosome of *E. coli MG1655 ΔlacZYA::Acatsac1/pSIM5tet* was performed by PCR with: 5’-AGGCGAAGCGGCATGCATTTACGTTGACACCATCGGCTTCTAGAGAACCCCTTG-3’ and 5’-ACTGGATCTATCAACAGGAGTCCAAGCCAAATTTATTATTAAGCACCGGTGGAG-3’; the binding parts on BioBrick prefix and mRFP1 ORF 3’ are underlined. The amplified fragments were integrated into the chromosome by lambda red recombineering (Datta *et al*., 2006) and sacB counter-selection (Gay *et al*., 1985). Integrations were confirmed by PCR and sequencing. The *E. coli* MG1655 genome-coded T7 RNAP mutant R632S was constructed by single-stranded DNA recombineering with mutagenic oligonucleotides and sucrose counter-selection by the Brandis lab (Gay *et al*., 1985; Ellis *et al*., 2001; Brandis *et al*., 2016). Briefly, the catsacB cassette was PCR amplified and replaced the C1894 of the T7 RNAP gene with the oligos: ΔC1894_fw: 5’-GGCTGGCTTACGGTGTTACTCGCAGTGTGACTAAGGTGTAGGCTGGAGCTGCTTC and ΔC1894_rv: 5’-CTCTTTGGACCCGTAAGCCAGCGTCATGACTGAACCATATGAATATCCTCCTTAGTTCC. The catsacB was then substituted with single nucleotide A using the ssDNA oligo 5’ GGCTGGCTTACGGTGTTACTCGCAGTGTGACTAAGAGTTCAGTCATGACGCTGGCTTACGGGTCCAAAGAG to mutate C1894 to A (Arg 632 to Ser). Point mutations of promoters on plasmids were made by inverse PCR (Hemsley *et al*., 1989).

### Quantitation of fluorescent protein expression

**For promoter assays**, *E. coli* MG1655 cells with chromosome integration or transformed with pSB3K3 or pSB1A2 plasmids (iGEM registry) harboring the mRFP1 expression cassettes were grown from the inoculation of the single colonies for 16 h at 37 °C in 1 mL LB media without or with 30 μg/mL kanamycin or 100 μg/mL ampicillin, respectively. Afterwards, 200 μL of the cells were transferred to the 96-well plates, and the fluorescence was measured by an Infinite M200 pro plate reader (Tecan) at 531/595 nm for mRFP1 with auto-gain. A_600_ was measured simultaneously. Fluorescence per cell was calculated by mRFP1 fluorescence/A_600_. All values were subtracted from the LB media with the respective antibiotics. The values were normalized to BBa_J23110 if needed. **For termination assays**, the mono- and bicistronic constructs were transformed into *E. coli* BL21 (DE3) for T7 RNAP or DH5α for *E. coli* RNAP termination assays, respectively. Single colonies were inoculated in 1 mL LB media supplemented with 30 μg/mL kanamycin, and the diluted overnight cultures were grown to mid-log phase (OD_600_ = 0.4–0.6) and subsequently induced with IPTG to relieve LacI-mediated repression. Cells were harvested after 4h with 1 mM IPTG induction for T7 and after 16h with 10 mM IPTG induction for *E. coli* RNAP owing to lower protein yields, respectively. The 200 μL harvested cells were treated with 33 μg/mL chloramphenicol at room temperature for 40 min to inhibit further protein expression and allow maturation of fluorescent proteins. Expression levels of mRFP1 and fwYellow were quantified by fluorescence measurement using the Tecan plate reader described above at 531/595 nm for mRFP1 with auto-gaining and 503/540 nm for fwYellow.

### Reverse transcription and sequencing

Total RNAs from the overnight-cultured *E. coli* DH5α cells expressing mRFP1 from different promoters on pSB1A2 high-copy plasmids were extracted and purified by ethanol precipitation (“RNAsnapTM RNA isolation method for Gram-negative bacteria” and “Sodium acetate/ethanol precipitation method” in Stead *et al*., 2012. Primer (Fig. 2, Integrated DNA Technologies) was gel purified, 20 pmol was kinased with 10 μCi γ^32^P-ATP (Revvity) and purified using Sephadex G50 (Cytiva). Reverse transcriptions were performed (Guo and Pyle, 2023) with minor modifications. Five μg of the total RNA, 40 μM of each dNTP, 70 μM of corresponding ddNTP and 5 pmol of the ^32^P-labelled primer were mixed in a 14.5 μL mixture, then 4 μL 5x RT buffer, 20 U RiboLock and 200 U Maxima Reverse Transcriptase (ThermoFisher) were added to a final volume of 20 μL. Mixtures were incubated at 50 °C for 30 min and heat-inactivated at 85 °C for 5 min. The samples were mixed with an equal volume of 2x formamide RNA loading dye (NEB) and incubated at 95 °C for 2 min, then cooled on ice for 5 min, and fully loaded onto a 12.5% polyacrylamide gel containing 8M urea. Visualization was by an Amersham Typhoon phosphor imager (Cytiva).

### Bioinformatic analysis of promoters

Source datasets were obtained from three independent studies:

1. Yona *et al*. (2018), Supplementary Data 1;
2. Lagator *et al*. (2022), https://github.com/szarma/Thermoters; and
3. Fuqua and Wagner (2026), https://github.com/tfuqua95/random-genomic.

Sequence files and corresponding activity measurements were imported into Microsoft Excel for preprocessing. All sequences were first screened for the presence of the TGTTA motif as the “docking sequence”. From this subset, sequences with predicted typical σ^70^ promoter features were removed using **SAPPHIRE** (https://sapphire.biw.kuleuven.be/, 83.3% accuracy; Coppens *et al*., 2022). The remaining sequences were further filtered using **MLDSPP (MG-D mode)** (https://mldspp-app-promoter.streamlit.app/, 98% accuracy; Paul *et al*., 2024) to exclude general bacterial promoters (*e*.*g*. Tab. S1). The resulting datasets, i.e. sequences containing the TGTTA motif but lacking typical bacterial promoter signatures, were then plotted based on their measured transcriptional activities. Sequences exhibiting activities above the respective detection thresholds defined in their original publications were classified as atypical promoters. For all identified atypical promoters, sequences were aligned relative to the TGTTA motif. Nucleotides upstream of this motif, spanning positions −42 to −16, were extracted for analysis. Heat maps were generated to quantify positional base biases that correlated with promoter activity using the following approach: for each position and base type (*e*.*g*. −42A), all promoter activities corresponding to sequences containing that base at that position were selected and averaged. A base at a given position was considered to correlate with higher promoter activity if its average activity exceeded by at least 1.5-fold the mean activity of the other three bases at the same position. The blue color scales are evenly distributed from the highest value (darkest) to the lowest (white), representing the correlation between base occurrence frequency and relative promoter activity at each position, with darker shades indicating bases that are preferentially enriched in higher-activity promoter sequences and *vice versa*. Sequence logos were generated with https://weblogo.berkeley.edu/logo.cgi, with default settings.

## ASSOCIATED CONTENT

### Data Availability Statement

The original data are available upon request.

### Supporting Information

Table S1

Figures S1 to S10

Graphical abstract

Supporting references

## AUTHOR INFORMATION

### Author contributions

L.B. and A.C.F. conceptualization and writing. L.B. methodology, investigation, formal analysis, data curation; A.C.F. supervision, project administration and funding acquisition.

## ACKNOWLEDGEMENTS

We are very grateful to Yiwen Zhang and Dr. Gerrit Brandis for reconstruction of the T7 RNAP mutant.

