## Supplementary material for "Transcription terminators with context-dependent promoter and terminator activities": Sup Tables and Figures

**CONTENTS**

Table S1

Figures S1 to S10

Graphical abstract

Supporting references

| **Sigma factor** | **-35 element** | **-10 element** | **Function** | **References** |
|---|---|---|---|---|
| $\sigma^{70}$ (RpoD) | TTGACA | TATAAT | housekeeping sigma factor | Lonetto *et al.*, 1992; Paget and Helmann, 2003 |
| $\sigma^{38}$ (RpoS) | TTGACA | TGTGCTATACT | stationary phase | Tanaka *et al.*, 1995; Schellhorn, 2020 |
| $\sigma^{32}$ (RpoH) | CTTGAA | CCCCATNT | heat shock | Erickson & Gross, 1989; Nonaka *et al.*, 2006 |
| $\sigma^{54}$ (RpoN) | CTGGCAC(-24) | TTGCA(-12) | nitrogen limitation | Hirschman *et al.*, 1985; Zhao *et al.*, 2010 |
| $\sigma^{28}$ (RpoF) | TAAA | GCCGATAA | flagellar genes | Chilcott & Hughes, 2000; Koo *et al.*, 2009 |
| $\sigma^{24}$ (RpoE) | GGAACTT | TCTGA | extreme heat stress | Raina *et al.*, 1995; Hiratsu *et al.*, 1995 |
| $\sigma^{19}$ (FecI) | GGAAAT | TGTCCT | ferric citrate transport | Enz *et al.*, 2003; Mahren & Braun, 2003 |

**Table S1.** The seven σ factors in *E. coli*. The -35 and -10 sequence elements and functions of the promoters in response to certain growth conditions are listed.

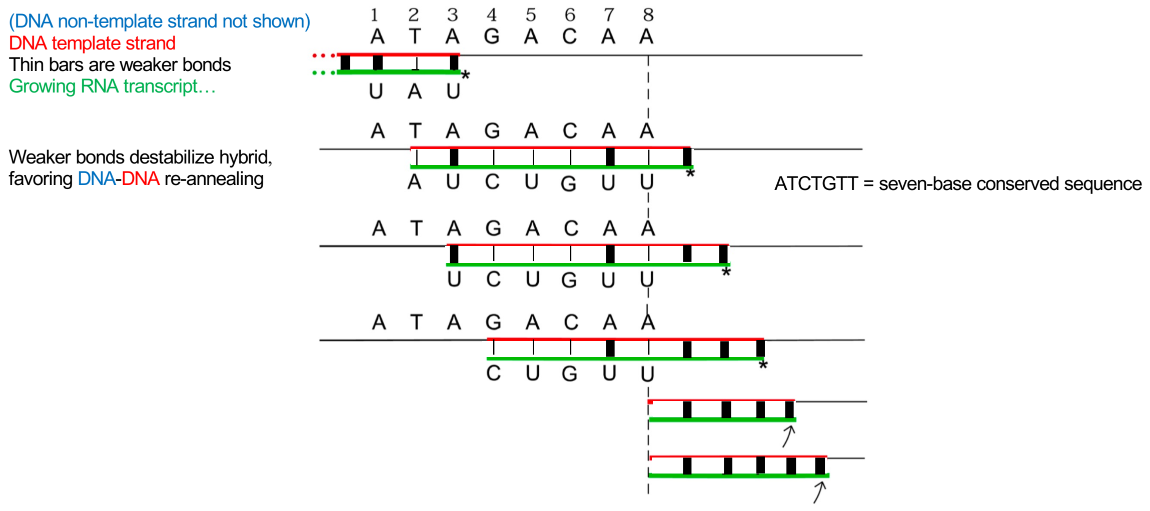


**Figure S1.** Model for termination by class II terminators based on base-pairing stabilities. A representation of progression of transcription through the CJ site is shown. Only the transcription hybrid formed between the DNA template strand (red) and the RNA (green) is shown. The vertical lines represent base pairs in this hybrid. Thinner vertical lines identify base pairs whose stabilities in the RNA-DNA hybrid are significantly lower than those in the DNA-DNA duplex. The asterisk denotes the last base that has been transcribed; the arrow denotes the base that is going to be transcribed. Modified from Zhang and Russu (2014).

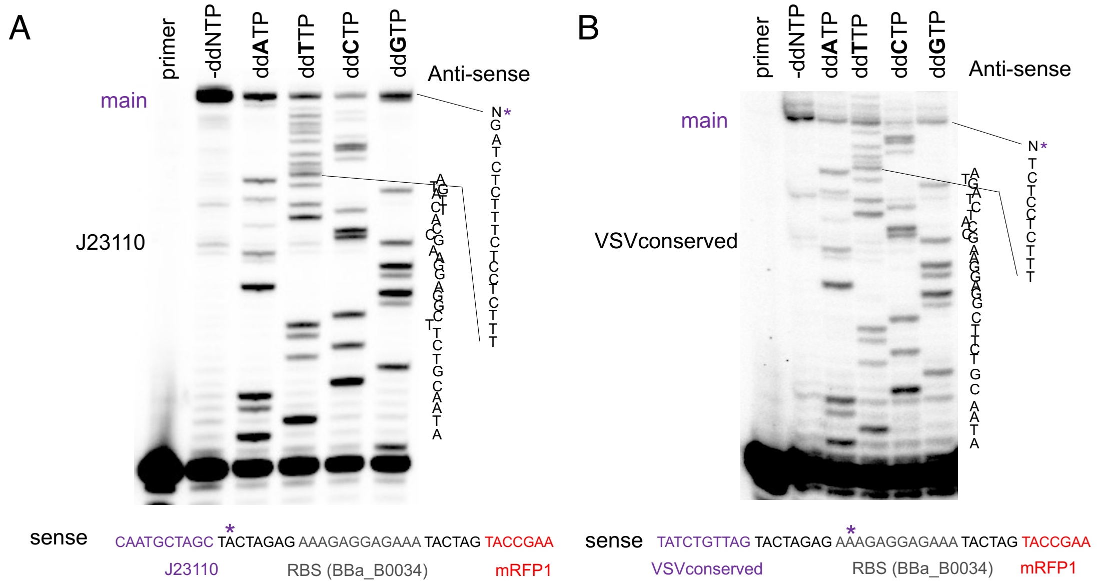


**Figure S2.** Mapping of transcription start sites of J23110 and VSVconserved promoter constructs. Representative gels are visualized by phosphorimaging and main stop sites are marked with asterisks. The whole gels are shown in Fig. S3.

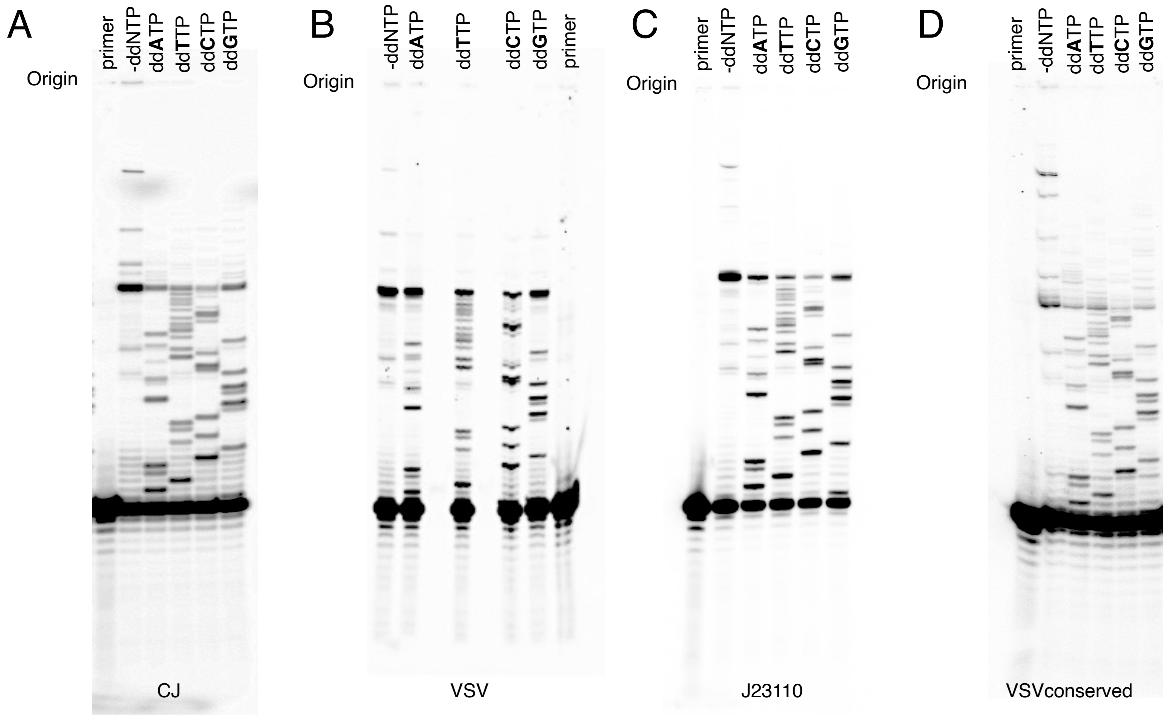


**Figure S3.** Whole gels from Fig. 2 and Fig. S2.

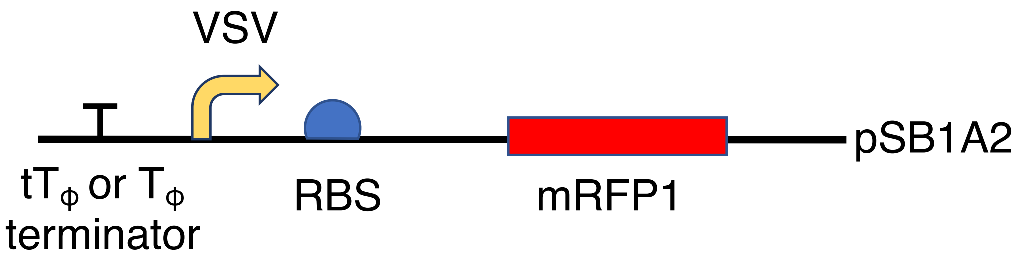


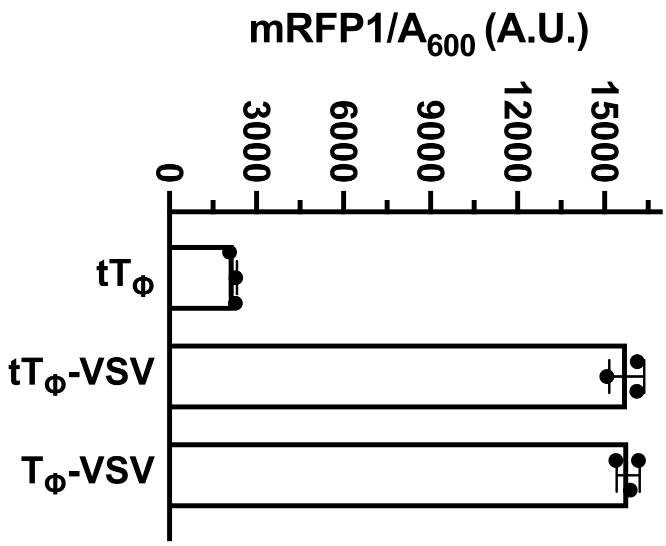


**Figure S4.** mRFP1 expression in overnight liquid cultures from VSV downstream of $tT_{\varphi}$ or $T_{\varphi}$. Error bars are standard deviations, n=3.

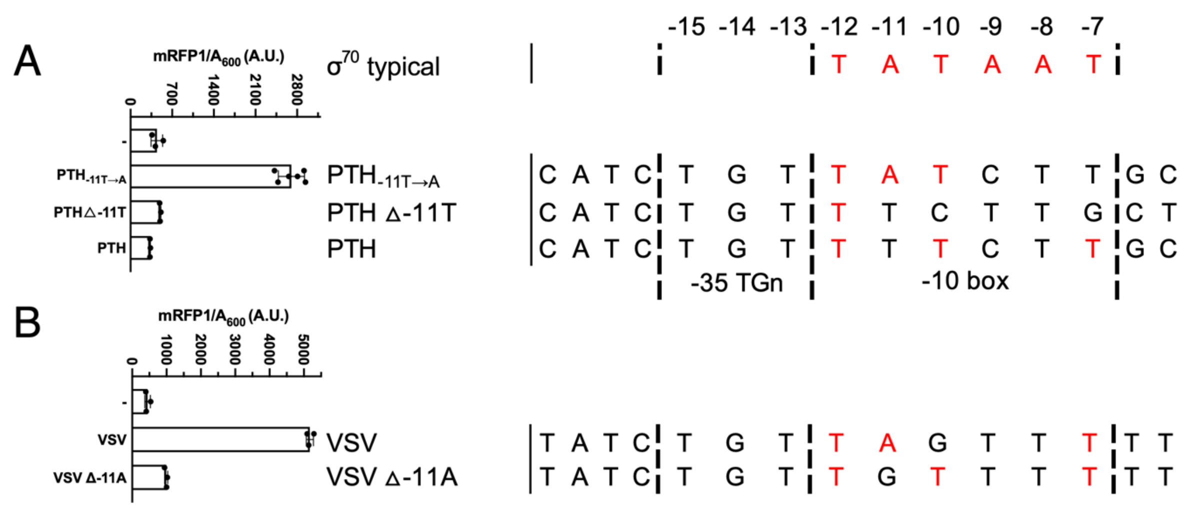


**Figure S5.** Effects of small deletions in the -10 boxes of PTH (A) and VSV (B) promoters on mRFP1 fluorescence at 37°C. Resulting base alignments that match the $\sigma^{70}$ -10 box consensus sequence are in red. Error bars are standard deviations, n=3.

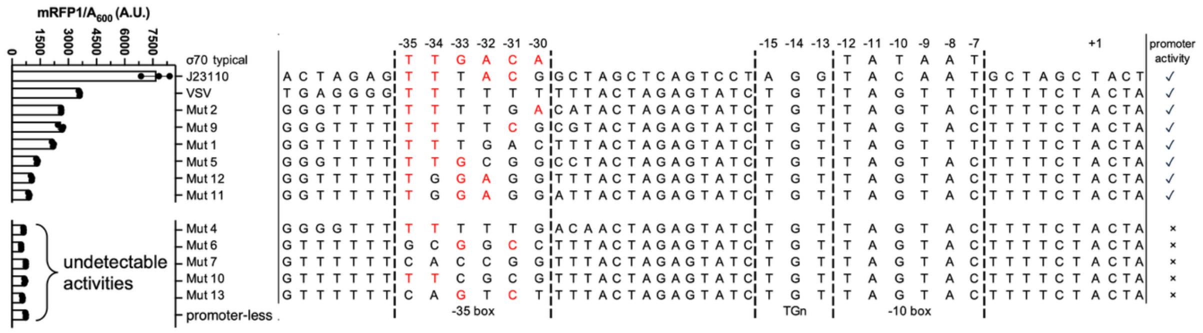


**Figure S6.** Effects of mutations around the -35 box of VSV on mRFP1 fluorescence at 37°C. Error bars are standard deviations, n≥3.

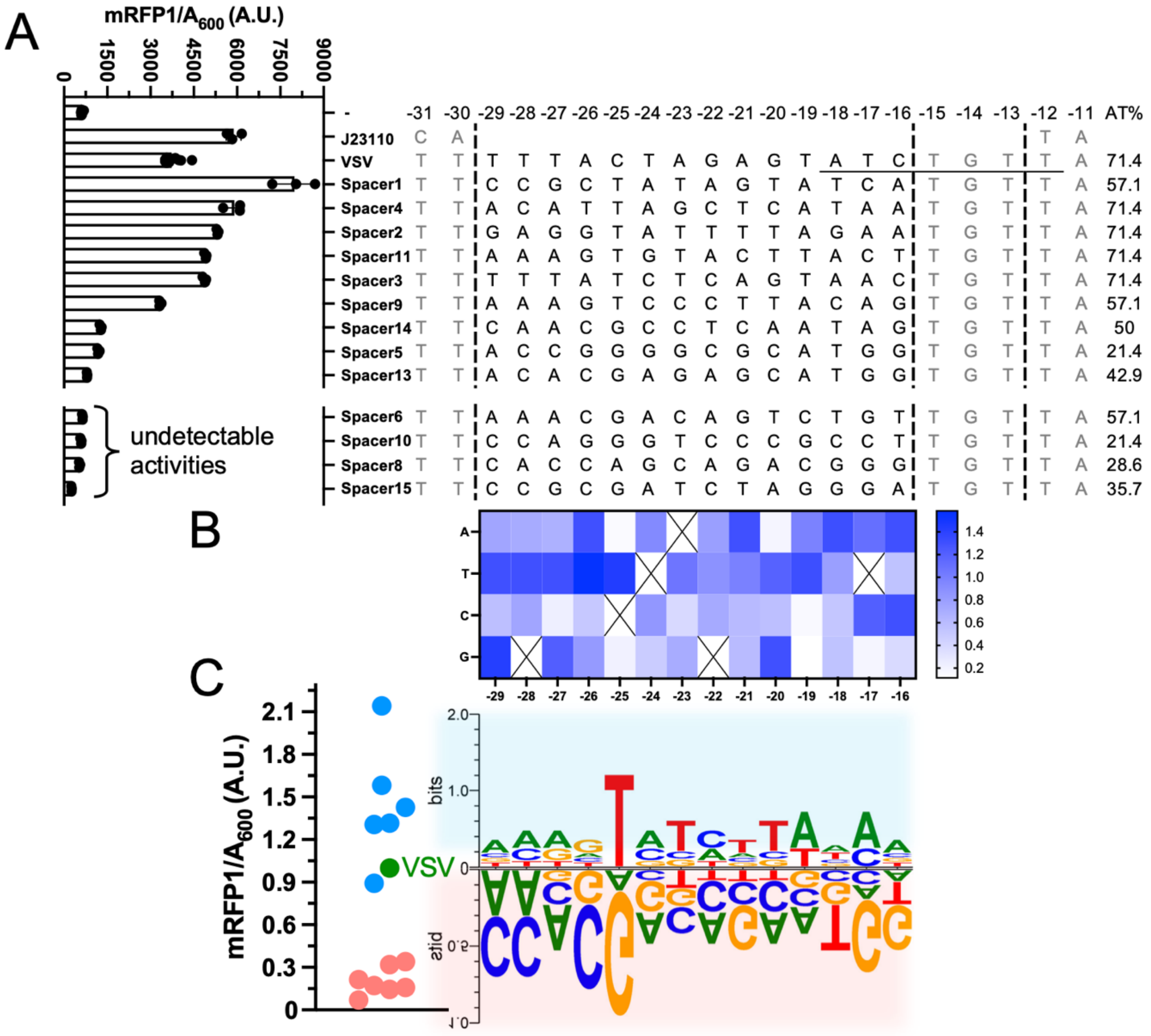


**Figure S7.** Effects of randomizing between the -35 and -10 boxes of VSV on mRFP1 fluorescence at 37°C. (A) mRFP1 fluorescence and sequence alignment of 13 randomized mutants. Conserved class II termination sequence is underlined. (B) Heatmap analysis showing base preferences across the randomized region. Promoter activities are normalized to VSV as 1.0. (C) Sequence logo analysis of activation (top) or inhibition (bottom) of the promoter activities. The sequence logo generated from red dots (inhibition) is shaded in red, and the other blue-dots sequence logo is shaded in blue. Error bars represent standard deviations (n = 3).

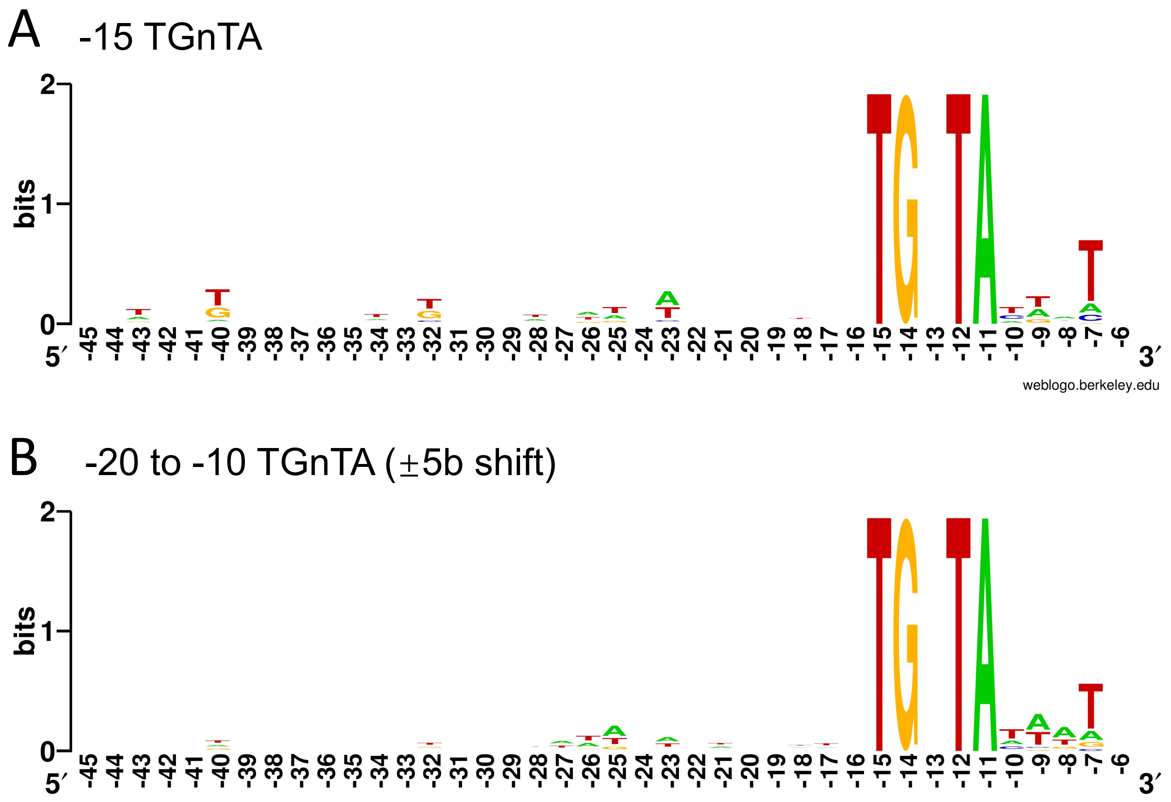


**Figure S8.** Sequence logo analysis of the *E. coli* orphan promoters of Thomason *et al.* (2015). TSSs not associated with known protein-coding genes (termed orphan TSSs) that harbored TGnTA starts either at the -15 position (A), or with a ±5 base shift (B) upstream of the TSSs, are plotted according to the base frequencies at each position.

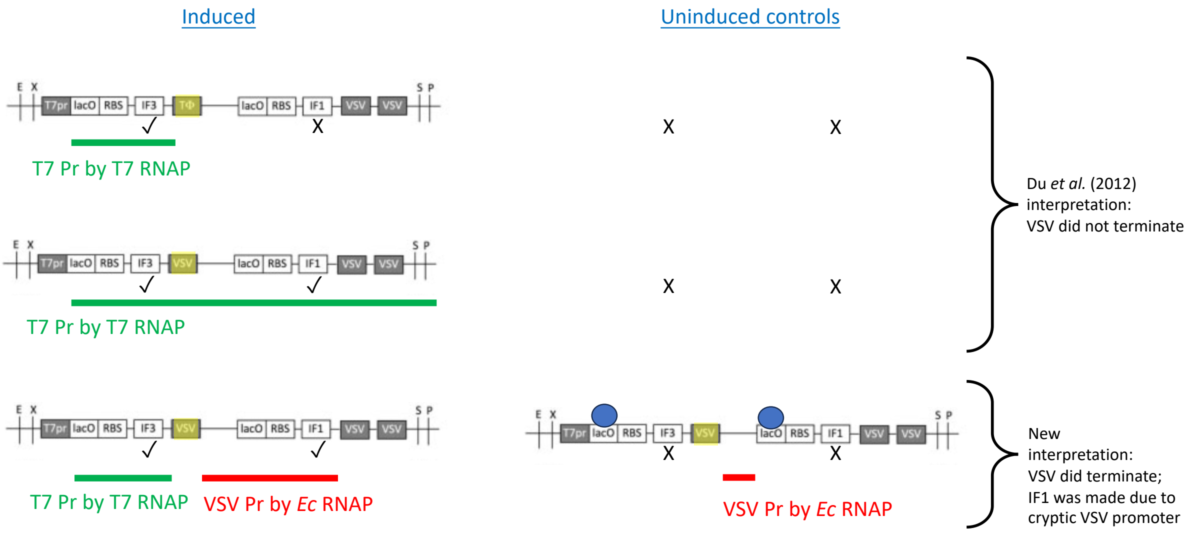


**Figure S9.** Opposite interpretations of results of an *in vivo* termination assay. The bicistrons were designed to produce IF3 protein +/- IF1 protein (dependent on strength of terminators shaded yellow) from an mRNA initiated from a single promoter for T7 RNAP upon induction with IPTG. Proteins produced by the two terminator constructs are indicated by ✓ and X (presence or absence of strong gel bands in Du *et al.,* 2012). Proposed transcripts are lines colored green (from T7 RNAP) and red (from *E. coli* RNAP). DNA-bound lac repressors are blue, illustrating how the undetectable background expression of IF1 under uninduced conditions was apparently due to blocked transcription initiation from VSV by repressor binding to the lacO operator upstream of IF1. Images of DNA constructs are modified from Du *et al.,* 2012.

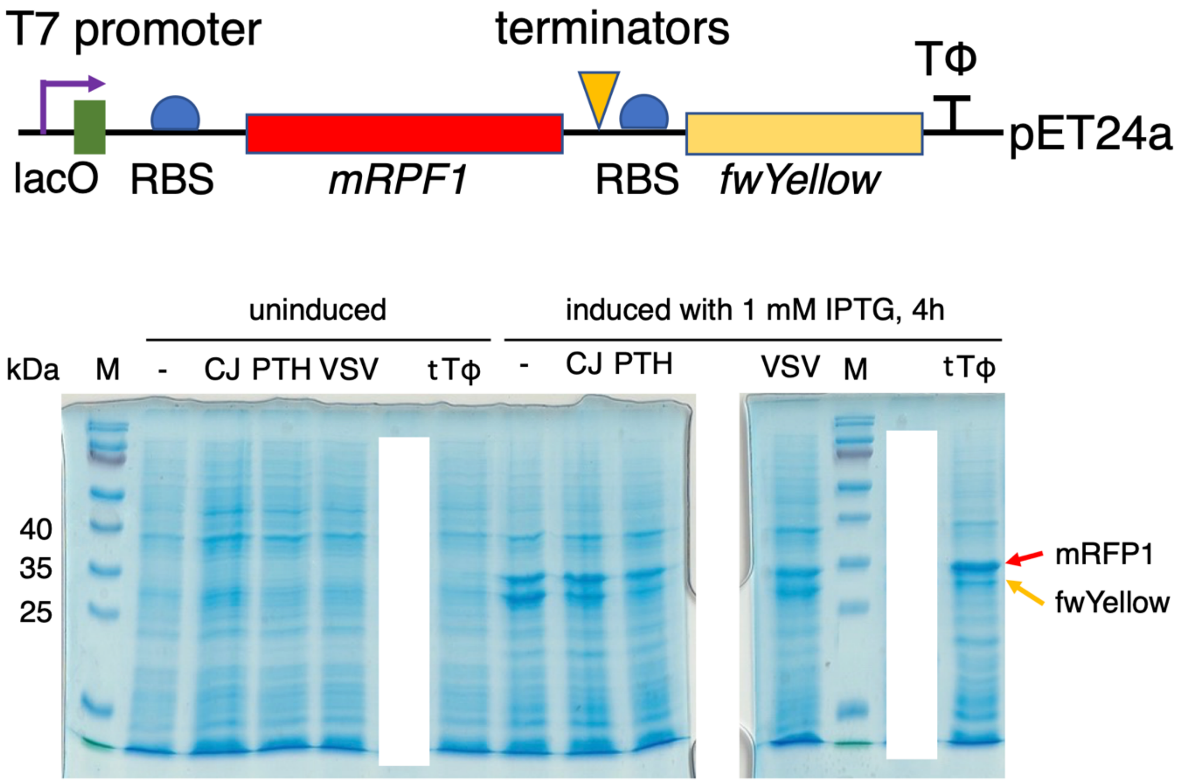


**Figure S10.** Representative SDS-15% PAGE of mRFP1 and fwYellow overexpressions with 1mM IPTG inductions for 4h from the bicistronic constructs from *E. coli* BL21 (DE3) cells. Denaturing gels stained with Coomassie Blue are total proteins from mixture of the three biological replicates from Fig. 6B after boiling.

Graphical abstract

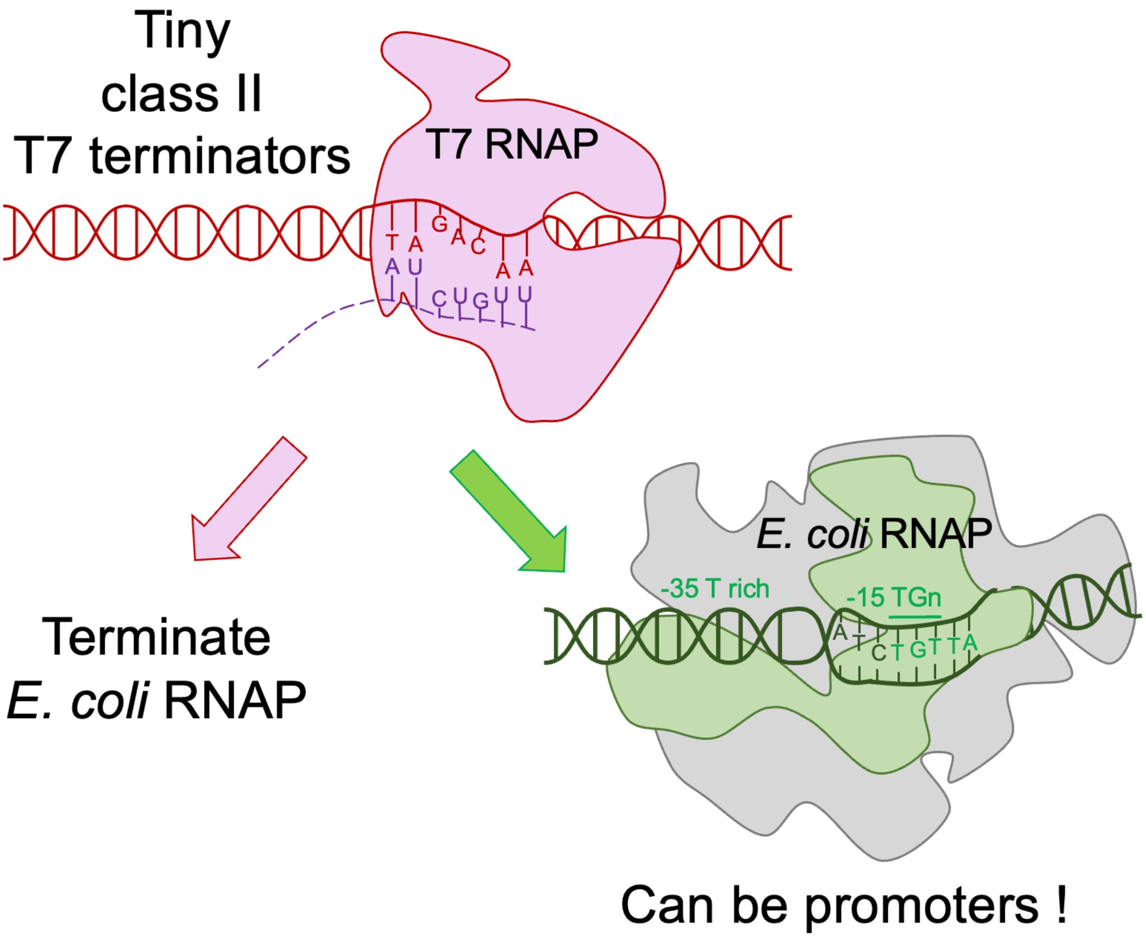


----
